# Proteasomal control of anti-CRISPRs for the regulation of CRISPR/Cas9 activity using Cas9-ACROBAT

**DOI:** 10.1101/2024.05.13.593596

**Authors:** Timothy D. Martin, Emma V. Watson, Mei Yuk Choi, Behnam Nabet, Nathanael S. Gray, Qikai Xu, Stephen J. Elledge

**Author notes:** Correspondence may be addressed to: Stephen J Elledge. Department of Pharmacology, University of Virginia School of Medicine, Charlottesville, VA, USA. Department of Systems Biology, University of Massachusetts Chan Medical School, Worcester, MA, USA.

## Abstract

Small molecule-mediated proteasomal degradation of proteins is a powerful tool for synthetic regulation of biological activity. To control Cas9 activity in cells, we engineered an anti-CRISPR protein, AcrIIA4, fused to a degradation (dTAG) or small molecule assisted shutoff (SMASh) tag. Co-expression of the tagged AcrIIA4 along with Cas9 and riboswitch-regulated sgRNAs enables precise tunable control of CRISPR activity by small molecule addition.

## INTRODUCTION

The facile nature of CRISPR/Cas9 technology has revolutionized the way researchers manipulate genomes for biological study. Since Cas9 is a nuclease that generates double-stranded breaks in genomic DNA, it is of the utmost importance to maintain precise control over its activity to mitigate off-target activity. To better control Cas9 activity, researchers have generated higher fidelity mutants of Cas9^1^, doxycycline and blue light-inducible Cas9^2,3,4^, split Cas9 variants^5^, amongst other inducible Cas9 systems. The discovery of a class of proteins named anti-CRISPRs (Acrs) provides another approach to limit and control Cas9 function^6^. These Acrs are small proteins naturally found in bacteriophage that bind to a Cas9/sgRNA complex and prevent its association with DNA thus providing the phage with a resistance mechanism to CRISPR-based bacterial immunity^7,8^.

We sought to engineer a naturally occurring Acr CRISPR inhibitor, AcrIIA4, with a protein tag that would enable control over Acr protein stability via the addition of small molecules. A recent degradation tag termed the dTAG was developed that involves the use of a conditional degron and an FKBP12^F36V^ variant fused to your protein of interest^9^. The dTAG-13 small molecule itself has 2 functional parts: one part binds the FKBP12^F36V^ domain while the other part interacts with the cereblon E3 ligase complex, effectively recruiting the protein degradation machinery to your protein of interest and leading to its destruction via the ubiquitin proteasome system (UPS)^9^.

In this study we fused a dTAG to the C-terminus of AcrIIA4, which constitutively inhibits the Cas9/sgRNA complex when expressed. Treatment with the dTAG-13 compound causes E3 ligase recruitment to AcrIIA4 and its destruction via the UPS, thus unleashing CRISPR/Cas9 activity. As another level of regulation, we engineered sgRNAs to contain a theophylline cleavable riboswitch. Theophylline addition leads to riboswitch activation which then self-cleaves freeing the sgRNA to bind to Cas9. Combining the conditional degron-tagged AcrIIA4 with riboswitch sgRNAs allows for tight control over CRISPR/Cas9 activity in human cells.

### Regulation of CRISPR/Cas9 cutting using a degron-controlled anti-CRISPR protein, AcrIIA4

To test our degron-controlled AcrIIA4 system (Fig. 1A), we generated a series of lentiviral CMV-driven bicistronic vectors which express the AcrIIA4-dTAG fusion and different *S. pyogenes* Cas9 derivatives for DNA cutting, activation (CRISPRa), or inhibition (CRISPRi) separated by a T2A peptide (Fig. S1A). We performed a dTAG-13 dose response treatment to examine the loss of AcrIIA4 expression in human colonic epithelial cells (HCECs) stably expressing the DNA-cutting vector diagrammed in Fig. S1A. Treatment with dTAG-13 caused a decrease in AcrIIA4 expression after 72 h with a maximal effect seen near 200 nM (Fig. 1B), the dose used for all remaining studies unless otherwise noted. We examined the kinetics of cutting after dTAG-13 treatment using a sgRNA targeting a repetitive telomere sequence and examining DNA damage via H2A.X phosphorylation. Increased phospho-H2A.X was seen in as little as one hour after dTAG-13 treatment (Fig. S1B). Next, we took GFP^+^ HCECs expressing the AcrIIA4-dTAG components with Cas9 and transduced them with an *AAVS1* control sgRNA or a sgRNA sequence targeting *GFP*^10^. Treatment with either DMSO control or dTAG-13 did not affect GFP expression in cells expressing the control sgRNA (Fig. 1C), while a dTAG-13 dose-dependent loss of GFP was observed in cells expressing the GFP-targeting sgRNA (Fig. 1C).

**Figure 1.**
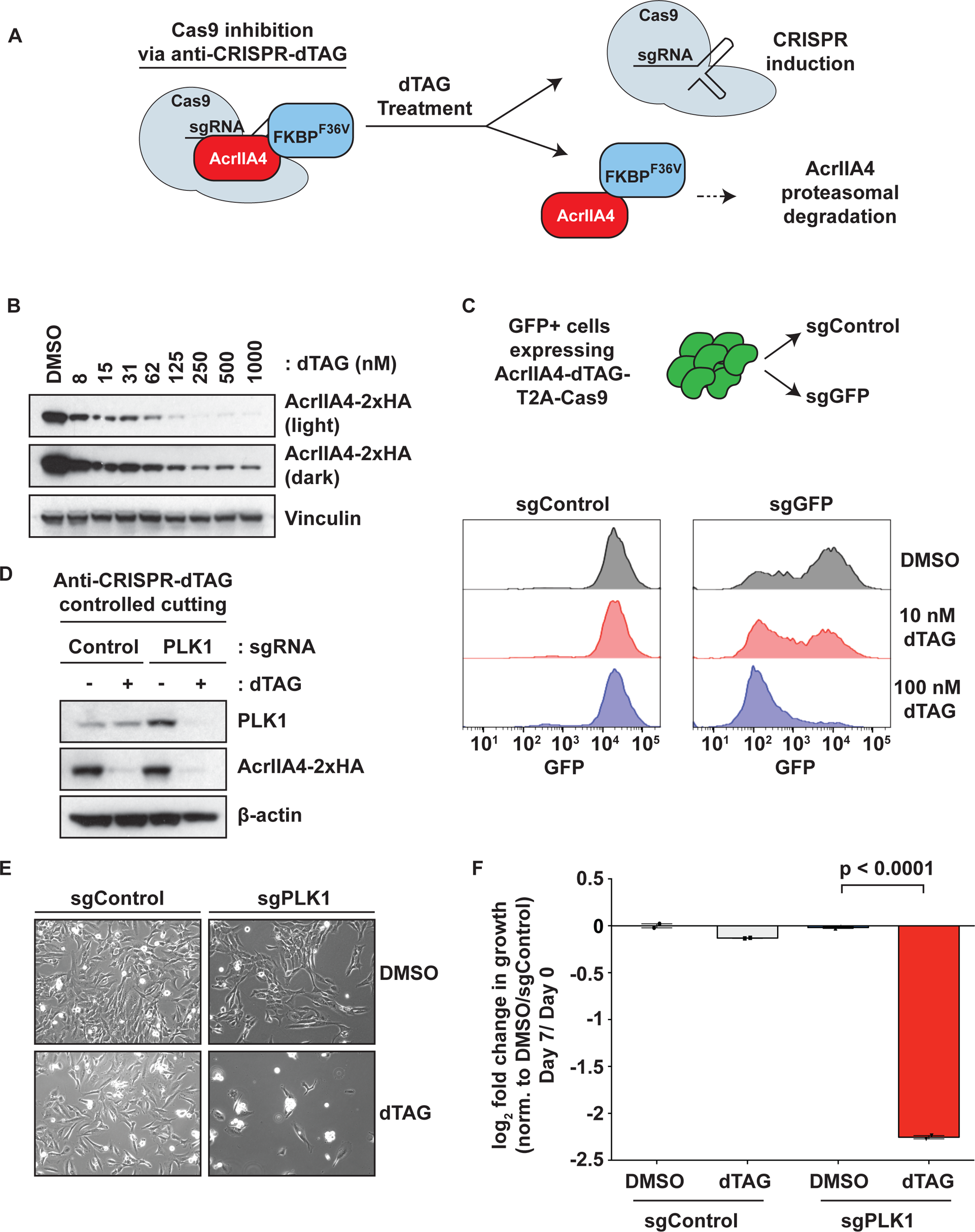
Regulation of CRISPR gene cutting via dTAG controlled AcrIIA4. **A.** The AcrIIA4-dTAG system. The dTAG fused anti-CRISPR protein AcrIIA4 binds to the Cas9/sgRNA complex preventing DNA binding. Treatment with dTAG-13 leads to AcrIIA4 proteasomal degradation unleashing CRISPR activity by allowing the Cas9/sgRNA complex to associate with DNA. **B.** Loss of AcrIIA4 expression via dTAG-13 dose response. HCECs stably expressing AcrIIA4-dTAG were treated with the indicated concentrations of dTAG-13 for 3 days. Extracts from these cells were immunoblotted for the indicated proteins to monitor AcrIIA4 loss. **C.** dTAG control of CRISPR cutting. HCECs stably expressing Cas9 with AcrIIA4-dTAG and GFP were transduced with sgRNAs targeting *GFP* or an *AAVS1* targeting control. Cells were treated with DMSO or the indicated concentration of dTAG-13 for 7 days. GFP fluorescence was determined by flow cytometry. **D.** dTAG control of endogenous gene cutting. HCECs stably expressing Cas9 with AcrIIA4-dTAG were transduced with sgRNAs targeting *PLK1* or an *AAVS1* targeting control. Extracts from these cells were immunoblotted for the indicated proteins after 5 days of treatment. **E.** AcrIIA4-dTAG controlled *PLK1* gene cutting causes cell death. Cells from Fig. 1D were imaged after 5 days of treatment with DMSO or 200 nM dTAG-13. **F.** Cutting of *PLK1* results in decreased cell proliferation. HCECs stably expressing Cas9 with AcrIIA4-dTAG and a control *AAVS1* or *PLK1* targeting sgRNA were treated with DMSO or 200 nM dTAG-13. Cells were counted after 7 days of treatment and the log_2_ fold change in cell number was determined. Fold change after 7 days is normalized to sgControl/DMSO.

To examine AcrIIA4-dTAG control of endogenous gene cutting we chose to target the essential mitotic gene *PLK1*. Since loss of PLK1 expression is lethal, this provides a readout of AcrIIA4-dTAG control of CRISPR activity via cell viability. HCECs stably expressing the AcrIIA4-dTAG and Cas9 construct were transduced with sgRNAs targeting *PLK1* or *AAVS1* (control). Treatment with dTAG-13 results in AcrIIA4 protein depletion that coincides with PLK1 protein loss in the cells expressing the *PLK1* sgRNA but not the AAVS1-targeting sgRNA control (Fig. 1D). As expected, activating CRISPR targeting of *PLK1* via dTAG-13 treatment increases cell death and reduces cellular proliferation and (Figs. 1E and 1F).

### Regulation of gene expression via CRISPRa and CRISPRi using a degron-tagged AcrIIA4

In addition to gene cutting and inactivation, CRISPR-regulated gene activation (CRISPRa) is a powerful tool for cell biology and genetic screening^11^. We used the AcrIIA4-dTAG system co-expressed with a nuclease-dead dCas9 fused to the <u>V</u>P64, <u>p</u>65/RelA, and <u>r</u>TA (VPR) transactivation domains to regulate the expression of a dTomato fluorescent reporter^12^. Human pancreatic nestin positive epithelial cells (HPNEs) stably expressing the AcrIIA4-dTAG with dCas9-VPR construct and a dTomato reporter were transduced with either *AAVS1* control or reporter-targeting sgRNAs. Treatment with dTAG-13 led to a dose-dependent increase in dTomato reporter expression that correlated with loss of AcrIIA4 protein expression (Fig. 2A). We next measured dTomato reporter expression using flow cytometry and found a dose dependent increase in dTomato expression (Fig. 2B). In these flow cytometry experiments we found that dTomato expression activation occurred in approximately 40 percent of cells (Fig. 2B). This is most likely due our use of a polyclonal population of cells transduced with three different constructs. Though cells were drug selected for construct integration, not all cells will contain equivalent and optimized expression of each component.

**Figure 2.**
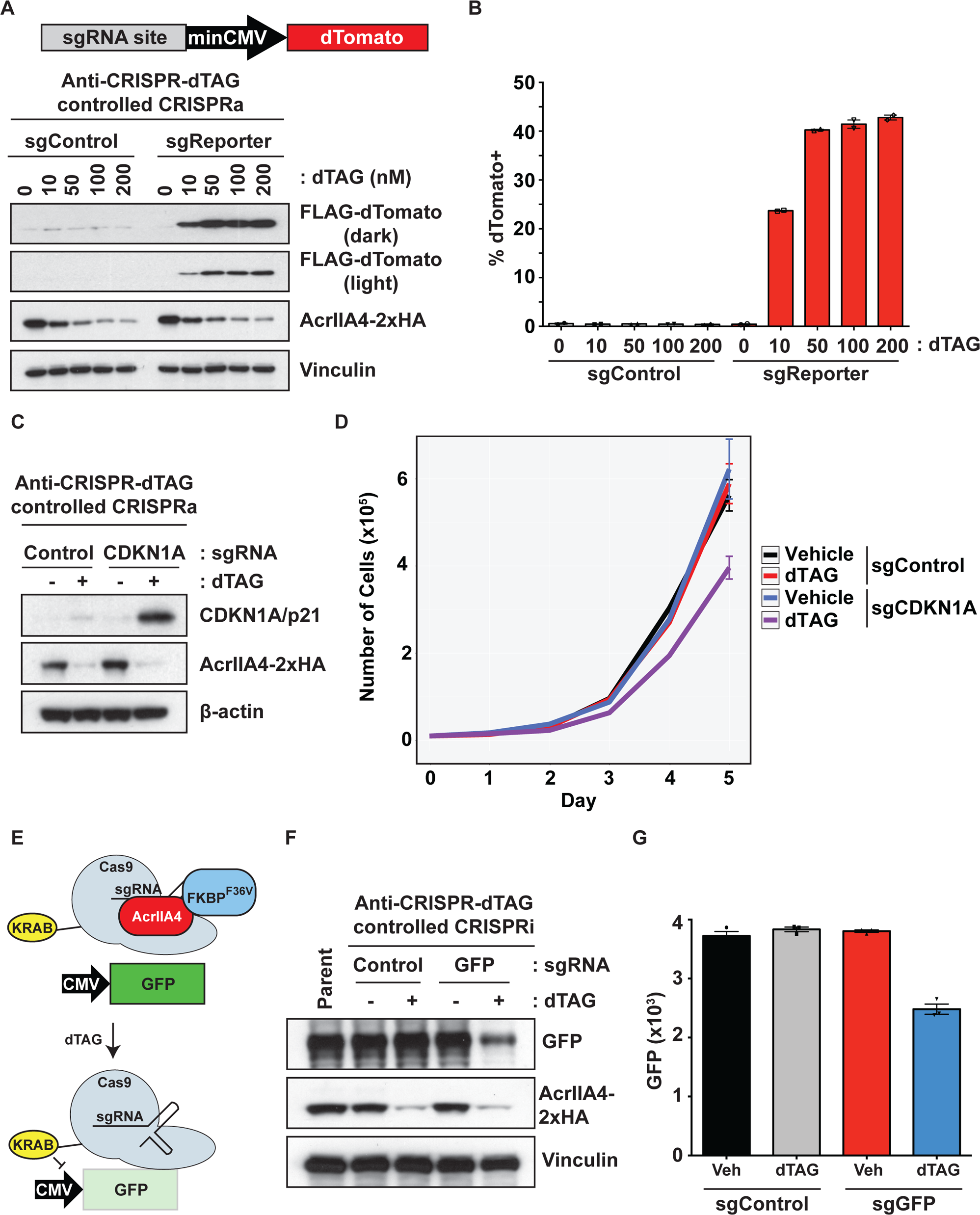
Regulation of gene activation via dTAG controlled AcrIIA4. **A.** Reporter gene activation controlled by AcrIIA4-dTAG. HPNEs stably expressing dCas9-VPR with AcrIIA4-dTAG, and a dTomato transgene controlled by a minimal CMV promoter were transduced with sgRNAs targeting the promoter (sgReporter) or an *AAVS1* targeting control. dTomato and AcrIIA4 expression were determined by western blot 5 days after treatment with DMSO or 200 nM dTAG-13. **B.** Reporter gene activation by AcrIIA4-dTAG controlled CRISPRa. Cells treated as in Fig. 2A were analyzed for dTomato expression by flow cytometry 5 days after treatment with DMSO or 200 nM dTAG-13. **C.** Endogenous gene activation via CRISPRa controlled by AcrIIA4-dTAG. HCECs stably expressing dCas9-VPR with AcrIIA4-dTAG were transduced with sgRNAs targeting *CDKN1A*/p21 or an *AAVS1* targeting control. Extracts from these cells were immunoblotted for the indicated proteins after 5 days after DMSO or 200 nM dTAG-13 treatment. **D.** Cell proliferation controlled by AcrIIA4-dTAG regulation of CRISPRa of *CDKN1A*/p21. Proliferation of cells from Fig. 2C after treatment with DMSO or 200 nM dTAG-13 was measured by cell counting over the indicated time period. **E.** Schematic of the anti-CRISPR control of CRISPRi. Treatment with dTAG-13 leads to destruction of AcrIIA4 and freeing of dCas9-KRAB to associate with the GFP reporter gene when a GFP sgRNA is present. This results in reduced GFP expression. **F.** CRISPRi control by AcrIIA4-dTAG. HCECs stably expressing GFP were transduced with the AcrIIA4-dTAG dCas9-KRAB construct shown in Fig. **S1C.** *AAVS1* Control or *GFP* targeting sgRNAs were expressed and cells were treated with DMSO or 200 nM dTAG-13 for 6 days. Extracts from these cells were immunoblotted for the indicated proteins to monitor loss of AcrIIA4 and reduction in GFP expression. **G.** Loss of GFP expression following dTAG-13 treatment. Cells from Fig. 2F were analyzed by flow cytometry to determine GFP fluorescence.

To test endogenous gene activation, we transduced HCECs stably expressing AcrIIA4-dTAG and dCas9-VPR with sgRNAs to activate *CDKN1A* or *AAVS1* (control). The *CDKN1A* gene encodes the p21 protein, a potent cell cycle regulator controlling cyclin dependent kinase (CDK) activity^13^. Cells expressing the *CDKN1A* but not the control sgRNA treated with dTAG-13 displayed an increase in p21 expression (Fig. 2C). As expected, this increase in p21 expression resulted in a decrease in cellular proliferation (Fig. 2D).

We tested the ability to control gene expression using CRISPR interference (CRISPRi) in GFP+ HCEC cells stably expressing the AcrIIA4-dTAG construct with nuclease-dead dCas9 fused to a transcriptional repressive KRAB domain (Fig. 2E). Cells were transduced with either *AAVS1* control or a *GFP* sgRNA. Treatment with dTAG-13 to degrade AcrIIA4 thus activating CRISPRi activity led to a decrease in GFP reporter expression only in cells expressing the GFP-targeting sgRNA as determined by western blotting (Fig. 2F) and flow cytometry (Fig. 2G).

### The SMASh-tag system as an orthogonal approach to controlling AcrIIA4 activity

We used the <u>S</u>mall <u>M</u>olecule <u>A</u>ssisted <u>Sh</u>utoff (SMASh) tag as an independent conditional degradation system to control AcrIIA4 protein expression^14^. In this approach a NS3 viral protease domain is fused to the amino terminus of AcrIIA4 preceded immediately by the NS3 protease substrate motif. Under basal conditions the NS3 protease cleaves itself from AcrIIA4 allowing for CRISPR inhibition. Treatment with the NS3 protease inhibitor asunaprevir prevents cleavage from AcrIIA4 leading to its proteasomal degradation and induction of CRISPR activity (Fig. 3A). We generated a similar set of lentiviral SMASh-tagged AcrIIA4 constructs to control CRISPR activity (Fig. S1C). Using HCECs stably expressing the SMASh-AcrIIA4 CRISPR cutting construct, we observed an asunaprevir dose-dependent loss of AcrIIA4 at 72 h and we used 1 μM for all additional experiments unless noted (Fig. 3B). We used GFP+ HCECs stably expressing SMASh-AcrIIA4 with Cas9 and transduced them with lentiviral sgRNAs targeting either *AAVS1* control or the *GFP* transgene. Following treatment with 1 μM of asunaprevir we observed a decrease in AcrIIA4 protein expression that correlated with a decrease in GFP expression (Fig. 3C). We took HPNE cells expressing the dTomato reporter described earlier in Fig. 2A and stably expressed a SMASh-AcrIIA4 construct that co-expressed dCas9-VPR to examine control of CRISPRa. Cells expressing a sgRNA targeting the reporter but not *AAVS1* control exhibited an asunaprevir dose dependent increase in dTomato reporter expression (Fig. 3D and 3E). Time course experiments revealed a peak in dTomato reporter expression following 5 days of continuous asunaprevir treatment (Fig. 3F and 3G).

**Figure 3.**
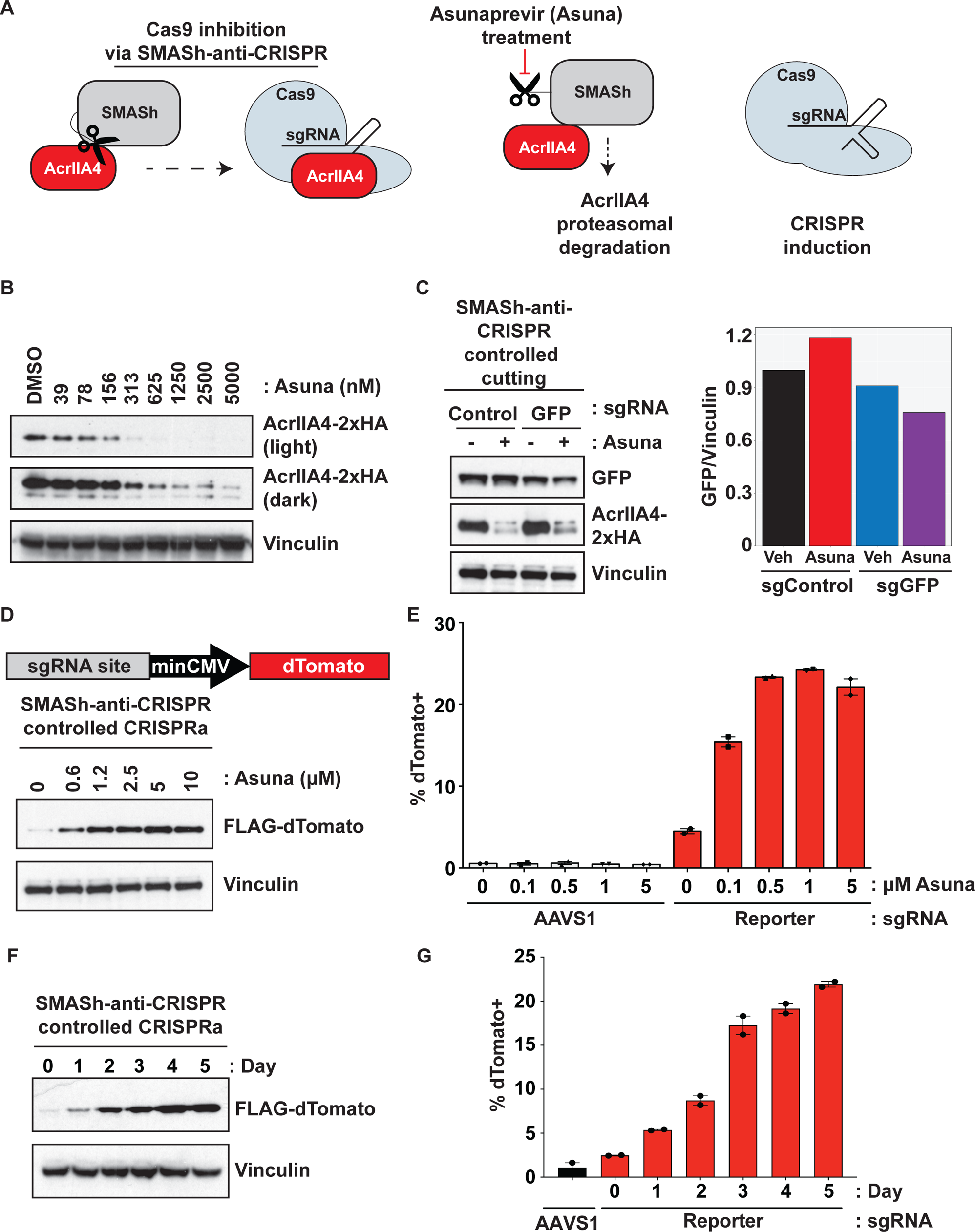
SMASh-tagged AcrIIA4 characterization. **A.** Schematic of SMASh-AcrIIA4 system. The SMASh-tagged AcrIIA4 binds to the Cas9/sgRNA complex preventing DNA binding following cleavage by the SMASh tag NS3 protease domain. Treatment with asunaprevir blocks NS3 protease activity which leads to AcrIIA4 proteasomal degradation. This activates Cas9 by allowing the Cas9/sgRNA complex to associate with DNA. **B.** Loss of AcrIIA4 expression via asunaprevir dose response. HCECs stably expressing SMASh-AcrIIA4 were treated with the indicated concentrations of asunaprevir for 3 days. Extracts from these cells were immunoblotted for the indicated proteins to monitor AcrIIA4 loss. **C.** Control of CRISPR cutting by SMASh-AcrIIA4. HCECs stably expressing Cas9 with SMASh-AcrIIA4 and GFP were transduced with sgRNAs targeting *GFP* or an *AAVS1* targeting control. Cells were treated with DMSO or 1 μM for 7 days. Cell lysates were western blotted with indicated antibodies to monitor loss of AcrIIA4 and GFP. Bar graph represents the ratio of GFP/Vinculin from the western blot. **D.** SMASh-AcrIIA4 and asunaprevir dose-dependent response of dTomato CRISPRa reporter via western blot. HCECs stably expressing dCas9-VPR with SMASh-AcrIIA4 and a dTomato transgene controlled by a minimal CMV promoter were transduced with sgRNAs targeting the promoter (sgReporter). Cells were treated with for 5 days with DMSO or the indicated concentrations of asunaprevir. Extracts from these cells were immunoblotted for the indicated proteins to monitor induction of FLAG-dTomato. **E.** SMASh-AcrIIa4 and asunaprevir dose-dependent response of a dTomato CRISPRa reporter. Cells from Fig. 3D were treated with DMSO or the indicated concentrations of asunaprevir for 5 days and were examined for dTomato fluorescence by flow cytometry. **F.** Time course of dTomato induction after treatment with asunaprevir. Cells described in Fig. 2D were treated with 1 μM of asunaprevir and harvested at the indicated time points. Cell lysates were immunoblotted with the indicated antibodies to monitor dTomato reporter induction. **G.** Time course of dTomato fluorescence after asunaprevir treatment. Cells from Fig. 3F were analyzed by flow cytometry for dTomato expression.

### Combining riboswitch regulated sgRNAs with degron-controlled AcrIIA4 for more precise control of CRISPR cutting activity

Our initial experiments using AcrIIA4-dTAG showed some leakiness in controlling CRISPR cutting activity as seen by the loss of GFP fluorescence even in the absence of dTAG-13 (Fig. 1C). To more tightly control CRISPR function we redesigned sgRNAs to include a small molecule-controlled riboswitch. We inserted an HDV ribozyme with an embedded theophylline riboswitch^15^ into the 3’ end of the guide RNA scaffold with a tail sequence complementary to a region spanning the tracrRNA stem-loop 1 and part of stem-loop 2. In the absence of theophylline, the riboswitch remains in the “off” position, with the stem-loop-blocking sequence preventing association of the guide RNA with Cas9. With addition of theophylline, the switch becomes active and self-cleaves to release the ribozyme/riboswitch and blocking sequence, enabling productive association of the guide RNA with Cas9. Importantly, this format enables universal control of sgRNAs independent of guide sequence. We have named the combination of degron controlled anti-CRISPR with riboswitch regulated sgRNAs the Cas9-ACROBAT system for <u>A</u>nti-<u>CR</u>ISPR <u>O</u>perator and <u>B</u>locker <u>A</u>t <u>T</u>racr (Fig. 4A).

**Figure 4.**
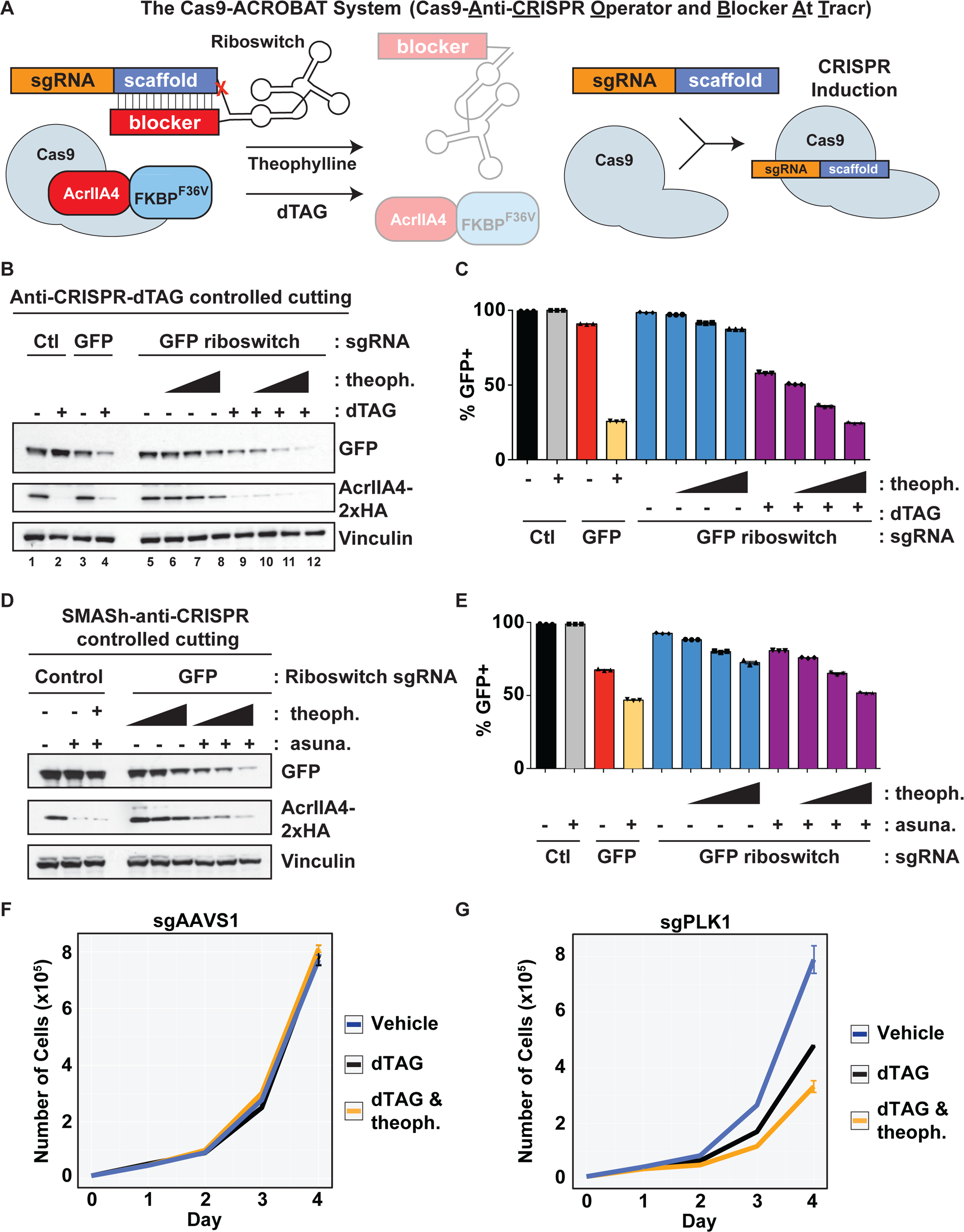
Riboswitch regulated sgRNAs combined with degron tagged AcrIIA4 to control CRISPR gene cutting activity. **A.** The Cas9-ACROBAT system. A theophylline riboswitch is added to the 3’ end of the sgRNA. Just 3’ to the riboswitch domain is a region of base complementarity to the sgRNA scaffold to hold it in an inactive conformation by preventing Cas9 binding. Treatment with theophylline causes riboswitch activation and cleavage of the riboswitch from the sgRNA freeing the sgRNA to bind Cas9. Combining the riboswitch sgRNAs with AcrIIA4-dTAG provides enhanced control over CRISPR activity. **B.** Riboswitch/dTAG control of CRISPR cutting. HCECs stably expressing Cas9, with AcrIIA4-dTAG and GFP were transduced with normal or riboswitch fused sgRNAs targeting *GFP* or an *AAVS1* targeting control sequence. Cells were treated with increasing amounts of theophylline (0, 0.1, 0.5, and 1 mM) and DMSO or 200 nM dTAG-13. Extracts from these cells were immunoblotted for the indicated proteins after 5 days of the indicated treatment. **C.** Loss of GFP fluorescence after dTAG/Riboswitch induction of Cas9 activity. Cells from Fig. 4B were examined for GFP fluorescence by flow cytometry. **D.** Repression of CRISPR activity via riboswitch and SMASh-AcrIIA4. HCECs stably expressing Cas9 with SMASh-AcrIIA4 and GFP were transduced with 19-nt blocking riboswitch fused sgRNAs targeting *GFP* or an *AAVS1* targeting control sequence. Cells were treated with increasing amounts of theophylline (0, 0.1, 0.5, and 1 mM) and DMSO or 1 μM asunaprevir for 7 days. Cell lysates were immunoblotted with the indicated antibodies to monitor loss of AcrIIA4 and GFP. **E.** Control of CRISPR gene cutting with SMASh-AcrIIA4 and riboswitch sgRNAs. HCECs stably expressing Cas9 with SMASh-AcrIIA4 and GFP were transduced with normal or 19-nt blocking riboswitch fused sgRNAs targeting *GFP* or an *AAVS1* targeting control sequence. Cells were treated as in Fig. 4D and GFP fluorescence was measured by flow cytometry 7 days after treatment. **F and G.** Riboswitch and dTAG control of endogenous gene cutting. HCECs stably expressing Cas9 with AcrIIA4-dTAG were transduced with riboswitch fused sgRNAs targeting an *AAVS1* control (panel F) or *PLK1* (panel G). Cells were treated as indicated (200 nM dTAG and 1 μM theophylline) and proliferation was measured by cell counting.

To test the Cas9-ACROBAT system we took HCECs expressing GFP with Cas9-ACROBAT for cutting and transduced them with traditional sgRNAs targeting *GFP* or *AAVS1* control, or with a riboswitch-regulated *GFP* sgRNA. Treatment with dTAG-13 results in a loss of AcrIIA4 expression independent of the sgRNA used (Fig. 4B). Importantly, treatment with 200 nM dTAG-13 and 1 mM theophylline in GFP riboswitch sgRNA cells resulted in equivalent loss of GFP fluorescence as seen with the use of traditional sgRNAs, indicating no loss in potency when using riboswitch sgRNAs (compare lanes 4 and 12 Fig. 4B) in the Cas9-ACROBAT system. As observed previously, the traditional sgRNA targeting GFP led to some leaky Cas9 activity even in the absence of dTAG-13 with loss of GFP in about 10% of cells (Fig. 4C). The Cas9-ACROBAT system, which includes the riboswitch-regulated sgRNA, eliminated this leakiness (Fig. 4C). Combined treatment of dTAG-13 with increasing amounts of theophylline led to an increasing loss of GFP fluorescence measured by flow cytometry (Fig. 4C).

We also engineered the riboswitch-regulated sgRNA without complementarity to the sgRNA scaffold and observed a very minimal loss of GFP expression upon theophylline addition indicating a necessity for base pairing between the sgRNA scaffold and riboswitch to properly inhibit CRISPR activity (Fig. S2A). Shortening the stem-loop base-pairing region from the 19-nt used in Fig. 4 to 16-nt also blunted the effect of the riboswitch sgRNAs at controlling CRISPR activity (Fig. S2B). For all subsequent experiments we used 19-nt complement riboswitches. In general, we found that the SMASh-tagged AcrIIA4 displayed leakier control of Cas9 activity than AcrIIA4-dTAG. To attempt to mitigate this leakiness, we combined SMASh-AcrIIA4 with riboswitch controlled sgRNAs from the Cas9-ACROBAT system and examined regulation of GFP expression. Treatment with theophylline and asunaprevir led to a dose dependent decrease in GFP expression that correlated with AcrIIA4 loss (Fig. 4D). Next, we compared traditional sgRNAs to riboswitch sgRNAs as in Fig. 4C. Traditional sgRNAs targeting *GFP* showed leaky Cas9 cutting activity in the absence of asunaprevir treatment which could be eliminated by using a riboswitch sgRNA paired with SMASh-AcrIIA4 (Fig. 4E). We found that both the complementarity and the length of the base pairing were critical for SMASh-tagged Cas9-ACROBAT similar to what we observed when using the dTAG version (Fig. S3A and S3B).

We tested endogenous gene cutting with Cas9-ACROBAT by targeting *PLK1* as done in Fig 1C. We monitored cell proliferation in HCECs expressing AcrIIA4-dTAG with Cas9 for gene cutting and riboswitch-regulated sgRNAs targeting control *AAVS1* or *PLK1*. Treatment with either dTAG-13 or dTAG-13 plus theophylline did not affect the growth of control *AAVS1* riboswitch sgRNA expressing cells (Fig. 4F). Conversely, we saw a decrease in proliferation in *PLK1* riboswitch sgRNA expressing cells after dTAG, which was further enhanced by theophylline treatment (Fig. 4G). Notably, untreated *PLK1* riboswitch sgRNA cells had a growth rate indistinguishable from control cells indicating strong suppression of CRISPR activity using the complete Cas9-ACROBAT system with riboswitch sgRNAs.

### The Cas9-ACROBAT system for regulating CRISPR activity

Lastly, we used Cas9-ACROBAT to control CRISPR gene activation. We used HCECs expressing dCas9-VPR with AcrIIA4-dTAG and a GFP reporter driven by a minimal promoter containing 4 copies of a sgRNA target site (Fig. 5A). Expression of a traditional sgRNA targeting a control sequence did not affect GFP expression but a traditional sgRNA targeting the GFP reporter led to robust GFP expression upon dTAG-13 treatment (Fig. 5A and 5B). Of note when using this GFP reporter system, even in the absence of dTAG-13 an increase of GFP is observed when a traditional sgRNA is used (compare lanes 1 and 3 Fig. 5A). This leakiness is completely eliminated when a riboswitch sgRNA is used instead (compare lane 3 and 5 in Fig. 5A).

**Figure 5.**
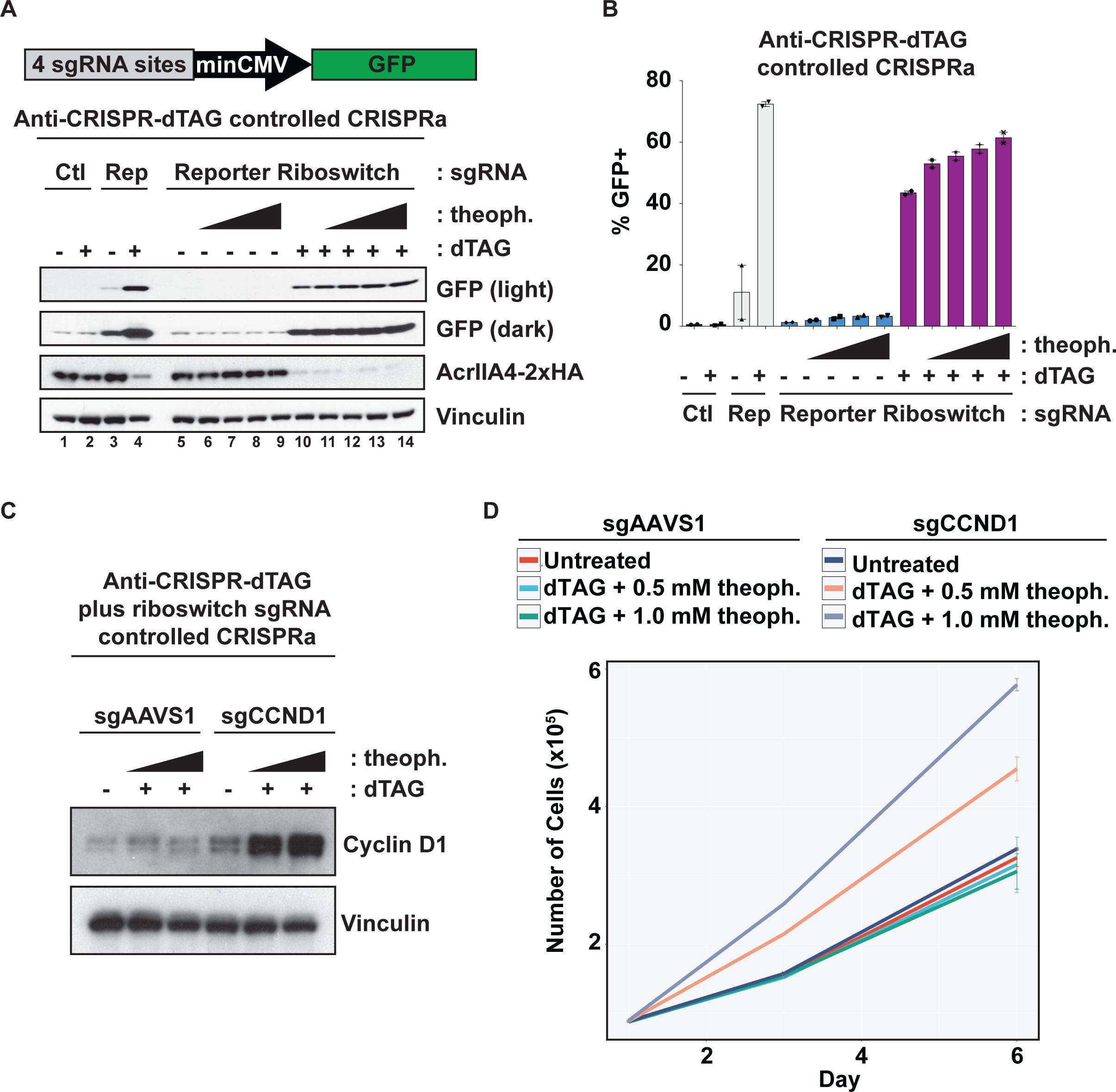
Riboswitch regulated sgRNAs and degron tagged AcrIIA4 to control CRISPR gene activation. **A.** Riboswitch and dTAG control of reporter gene activation. HCECs stably expressing dCas9-VPR with AcrIIA4-dTAG and a GFP transgene controlled by a minimal synthetic (SCP2) promoter were transduced with normal or riboswitch fused sgRNAs targeting the promoter (Reporter) or an *AAVS1* targeting control (Ctl). Cells were treated with increasing amounts of theophylline (0, 0.1, 0.5, and 1 mM) and DMSO with and without 200 nM dTAG-13. Extracts from these cells were immunoblotted for the indicated proteins after 5 days of the indicated treatment. **B.** Cells from Fig. 5A were examined for GFP fluorescence by flow cytometry. **C.** Riboswitch and dTAG control of endogenous gene activation. HCECs stably expressing dCas9-VPR with AcrIIA4-dTAG were transduced with riboswitch sgRNAs targeting control *AAVS1* or *CCND1/*cyclin D1. Cells were treated with theophylline (0, 0.5, and 1 mM) and DMSO with and without 200 nM dTAG-13. Extracts from these cells were immunoblotted for the indicated proteins after 5 days of the indicated treatment. **D.** Riboswitch regulation of CRISPRa of *CCND1*/cyclin D1 controls proliferation. Cells as described and treated in Fig. 5C were plated and counted over the indicated time course to monitor proliferation.

We used the Cas9-ACROBAT system to activate the endogenous expression of the gene *CCND1* which encodes the cyclin D1 protein, a potent driver of S phase entry and cellular proliferation. Expression of a riboswitch controlled sgRNA targeting *CCND1* but not *AAVS1* control increased the expression of cyclin D1 after dTAG and theophylline treatment (Fig. 5C). We measured proliferation of these cells and found a theophylline dose and dTAG dependent increase in growth when cyclin D1 expression was activated using Cas9-ACROBAT (Fig. 5D).

SMASh-tagged AcrIIA4 with dCas9-VPR coupled with the GFP reporter system from Fig. 5A showed substantial leakiness with the use of traditional sgRNAs (Fig. S3C). This leakiness was completely resolved without sacrificing potency by using Cas9-ACROBAT (Fig. S3C).

## DISCUSSION

Precisely controlling CRISPR/Cas9 activity is critical to avoid off-target activity. The discovery of the phage anti-CRISPR proteins provides a new way to limit the function of Cas9 to better control CRISPR experiments especially for future gene therapy procedures. Studies have shown control of CRISPR/Cas9 activity through regulation of AcrIIA4 by both light activation and a shield1-based system where destabilized AcrIIA4 becomes stable after addition of the small molecule shield1^16^. Each of these systems has their own advantages and drawbacks. Using light allows control over which specific cells in a population or even subcellular localizations within a cell receive protein activation but light activation is not an available technique to most labs and will have limited use for in vivo models depending upon which tissue needs light exposure. The destabilized version of AcrIIA4 requires continuous addition of shield1 to generate anti-CRISPR activity which has the potential for toxicity in cell culture and in vivo systems. Conversely, our Cas-ACROBAT system provides anti-CRISPR activity without the need for continuous small molecule treatment and provides the control of when to shut off AcrIIA4 function to induce CRISPR activity.

Work in *E. coli* demonstrated that riboswitch regulated sgRNAs are capable of controlling CRISPR function to increase transformation and editing efficiency allowing for the generation of combinatorial gene edits^17^. There have also been attempts to control CRISPR in HEK293T cells with theophylline activated sgRNAs but these have only been successful in controlling CRISPRi and not gene editing^18^. In our experiments we used the same concentrations of theophylline and were able to see CRISPR gene editing and CRISPRa induction. One possibility for this difference could be due to our cell line, HCECs, having better uptake of theophylline compared to HEK293Ts which may allow for better sgRNA cleavage. We show that combining our degron-fused Cas9 proteins with riboswitch regulated sgRNAs in human cells we are able to limit CRISPR cutting activity up to 95-98% using either the dTAG or SMASh tag degron system (Fig. 4B-4E) and prevent premature CRISPRa induction (Fig. 5A and 5B) without sacrificing potency.

Cas9-ACROBAT is an easy, user-friendly tool to aid researchers in controlling gene expression via CRISPR/Cas9. The Cas9-ACROBAT system provides more precise control over CRISPR activity and is an important addition to the ever-growing CRISPR toolbox.

## Supporting information

Supplemental DNA information

Supplemental spreadsheet of sgRNA sequences

Supplemental DNA sequence files

## ACKNOWLEDGEMENTS

We would like to thank all members of the Elledge lab for helpful feedback and discussions. T.D.M is supported by a Damon Runyon Cancer Research Foundation fellowship (DRC 2199-14). E.V.W is supported by a Damon Runyon Cancer Research Foundation fellowship (DRG-2269-16). B.N. is supported by an American Cancer Society Postdoctoral Fellowship (PF-17-010-01-CDD). This work was supported by the Katherine L. and Steven C. Pinard Research Fund to B.N. and N.S.G. and a grant from the NCI R01CA234600, the Cancer UK Grand Challenges Award and the Mark Foundation for Cancer Research to S.J.E. S.J.E is an investigator with the Howard Hughes Medical Institute.

## AUTHOR CONTRIBUTIONS

T.D.M, E.V.W, and S.J.E conceived of the study and designed experiments. T.D.M and M.Y.C performed all experiments. E.V.W designed and cloned the riboswitch/ribozyme-controlled sgRNA construct. Q.X. designed and cloned the GFP CRISPR activation reporter. B.N. and N.S.G provided reagents. T.D.M, E.V.W, and S.J.E wrote the manuscript with support from all other authors.

## ETHICS DECLARATIONS

B.N. is an inventor on patent applications related to the dTAG system (WO/2017/024318, WO/2017/024319, WO/2018/148443, WO/2018/148440). N.S.G. is a founder, science advisory board member (SAB) and equity holder in Syros, C4, Allorion, Lighthorse, Voronoi, Inception, Matchpoint, Cobro Ventures, GSK, Larkspur (board member), Shenandoah (board member), and Soltego (board member). The Gray lab receives or has received research funding from Novartis, Takeda, Astellas, Taiho, Jansen, Kinogen, Arbella, Deerfield, Springworks, Interline and Sanofi. S.J.E. is a founder and serves on the scientific advisory board of TScan Therapeutics, Immunity Bio and MAZE Therapeutics. S.J.E. is an Investigator with the Howard Hughes Medical Institute.

## METHODS

### Cell lines and culture

Human colonic epithelial cells (HCECs) were obtained from Dr. Jerry Shay (UT-Southwestern) and were cultured as described^19^. Human pancreatic nestin positive epithelial cells (HPNEs) have been previously described^20^. HEK293Ts were obtained from ATCC, grown in DMEM with 10% fetal calf serum (HyClone) and 1% penicillin/streptomycin (Gibco) and were used to generate lentivirus with pMD2.G and psPax2 second generation packaging vectors. All cells were tested for mycoplasma prior to use (Lonza, cat. # LT07-218). Proliferation was measured seeding 1,000 cells per well in a 96-well plate with CellTiter Blue (Promega,cat. # G8081) over the indicated time period and a plate reader (Perkin Elmer Victor X5, cat. #2030) with the Alamar blue setting or by direct cell counting with a Z Series Coulter Counter (Beckman-Coulter) over the indicated time period after initially plating 50,000 cells per well of a 12-well plate.

### Plasmids and other reagents

The following cDNAs were obtained from Addgene and the corresponding plasmids: AcrIIA4 (pJH376,plasmid #86842), T2A-spCas9 (pRubiG-T2A-Cas9, plasmid # 75348), dCas9-VPR (SP-dCas9-VPR, plasmid #63798), dCas9-KRAB (pLV-dCas9-KRAB-PGK-HygR, plasmid #83890), the dTomato reporter system (M-tdTom-SP, plasmid #48677), amino terminal SMASh-tag (pCS6-SMASh-YFP, plasmid #68852). AcrIIA4 was cloned into pENTR dTOPO and SMASh or dTAG domains were inserted via Gibson assembly (Clontech, cat. # 638909) to be in frame with AcrIIA4. The different Cas9 variants separated by a T2A sequence were next cloned into the SMASh or dTAG fused AcrIIA4 via Gibson assembly. Full AcrIIA4 and Cas9 constructs were then moved to pHAGE CMV DEST hygro via LR clonase (Invitrogen, cat. # 11791-020). Riboswitch/ribozyme sequences were previously described^15^. dTAG-13 was synthesized as previously described^21^.

All plasmid maps and sequences can be found as supplemental information as Supplemental DNA sequence files and reagents will be deposited with Addgene for distribution.

### Western blotting and antibodies

All cells were lysed in 1x RIPA lysis buffer (50 mM Tris-HCl, 150 mM NaCl, 1% NP-40, 0.5% sodium deoxycholate and 0.1% SDS, Boston Bioproducts, cat. # BP-115X) with protease and phosphatase inhibitors (Thermo, cat. # 78440). Protein was quantified by Bradford assay (Bio-Rad, cat. # 500-0006EDU). Protein lysates were mixed with 4X reducing sample buffer (Invitrogen, cat. # NP0007), separated on 4-12% Bis-Tris gels (Thermo, cat. # NP0336BOX), and transferred via a Trans-Blot Turbo Transfer System (Bio-Rad, cat. # 1704150) to nitrocellulose membranes (Bio-Rad, cat.# 170-4158). Membranes were blocked for 1 h at room temperature with 5% milk in TBST (Santa Cruz, cat. # sc-362311) and incubated with the following antibodies at the indicated dilutions in 5% BSA (VWR, cat. # VWRV0332) in TBST either overnight at 4C or 1 h at room temperature: Vinculin (1:10,000, cat. # V9131), HA-HRP (1:1,000, cat. # 12013819001) from Sigma, PLK1 (1:1,000, cat. # 4513T), β-actin (1:10,000, cat. # 3700S), GFP (1:1,000, cat. # 2956S), Phospho-Histone H2A.X Ser139 (1:1,000, cat. # 9718S), and FLAG-HRP (1:1,000, cat. # 2044S) from Cell Signaling, CDKN1A/p21 (1:1,000, cat. # OP64) from Millipore, and GAPDH (1:10,000, cat. # sc-365062) from Santa Cruz. Goat anti-mouse or rabbit HRP conjugated secondary antibodies (1:5,000, cat. #s 31430 and 31460) incubated for 1 h at room temperature were used for detection with ECL (Perkin-Elmer, cat. # NEL104001EA) via LI-COR (Odyssey model #2800) or film (HyBlot CL, Denville Scientific, cat. # 1159M38).

### Flow cytometry

Cells were harvested by trypsinization and filtered via a 5 mL FACs tube with 0.35 um cell strainer (Falcon, cat. # 352235) to ensure single cell suspensions. Samples were analyzed on a LSR II flow cytometer (Becton-Dickinson). Live, single cells used in the analyses were gated via SSC-A and FSC-A and FSC-H and FSC-A, respectively. GFP was detected using a 488 nm laser and FITC filter and dTomato was detected with a 568 nM laser and PE filter.

### Statistical Analyses

Differences between two groups were determined using an unpaired, two-tailed Student’s t test in Prism (Graphpad). Error bars denote S.E.M. All assays were performed in duplicate or triplicate as indicated and repeated for a total of at least two independent experiments.

---

**Figure S1.**
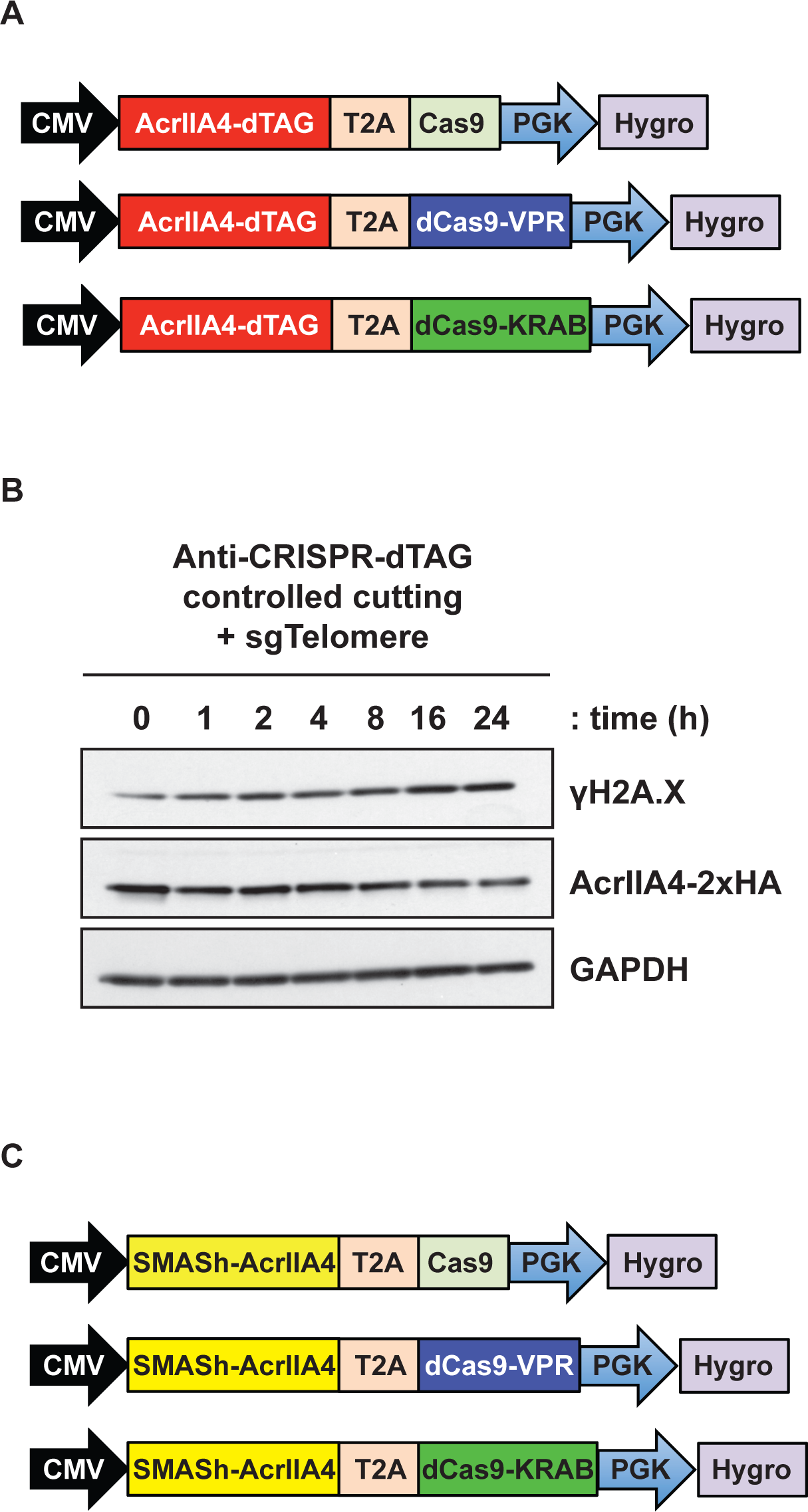
AcrIIA4-dTAG system and characterization. **A.** Lentiviral expression constructs for expression of dTAG components for gene cutting, CRISPRa, and CRISPRi. **B.** Time course of telomere cutting after dTAG-13 treatment. HCECs stably expressing Cas9 with AcrIIA4-dTAG and a telomere targeting sgRNA were treated with 200 nM dTAG-13 for the indicated time. Extracts from these cells were immunoblotted for the indicated proteins to monitor DNA damage due to telomere cutting via CRISPR induction. **C.** Lentiviral expression constructs for expression of SMASh-tagged components for gene cutting, CRISPRa, and CRISPRi.

**Figure S2.**
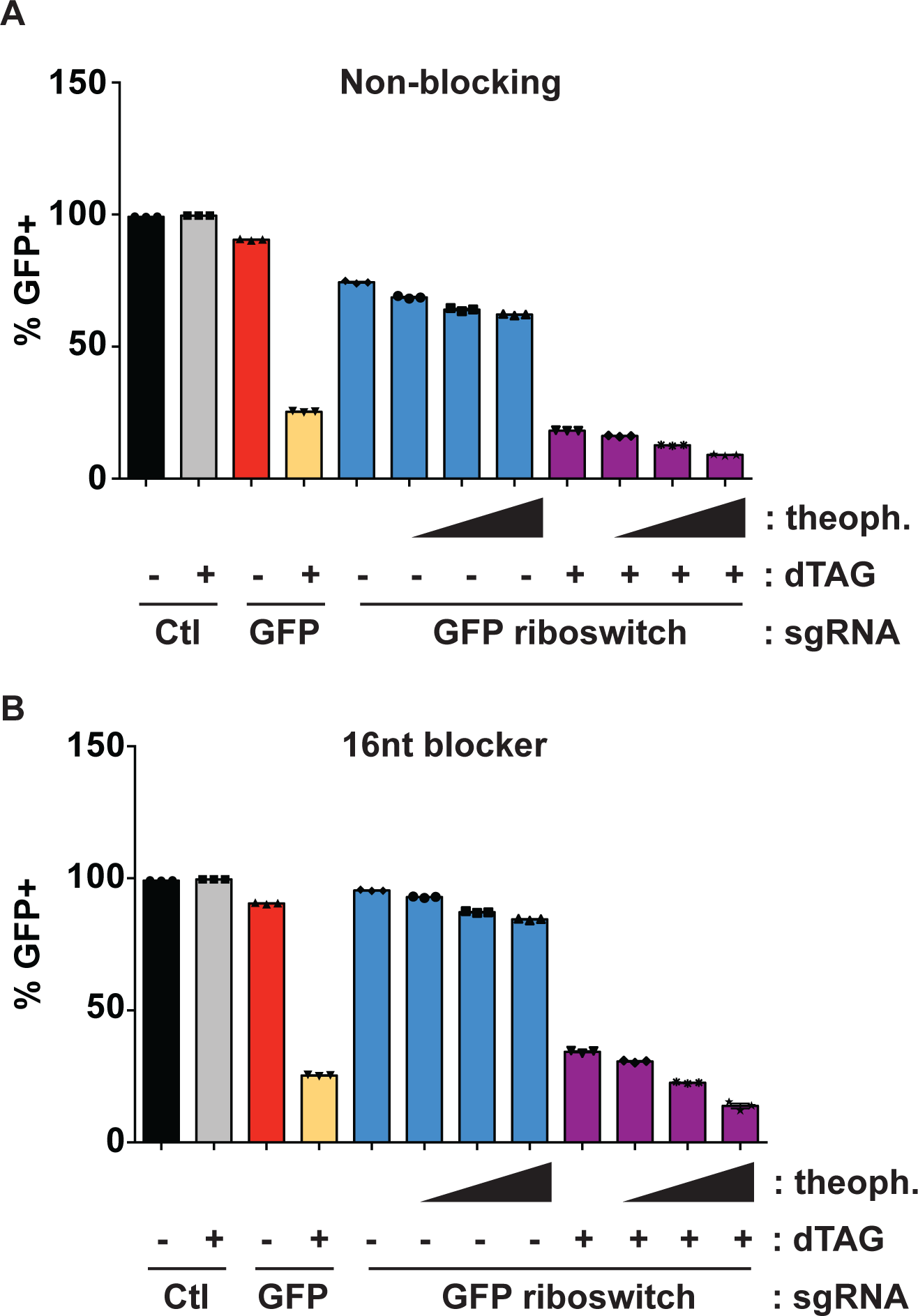
Comparison of non-blocking and 16-nt blocking riboswitches on GFP cutting controlled by AcrIIA4-dTAG. **A.** A non-blocking riboswitch does not repress CRISPR activity. HCECs stably expressing Cas9 with AcrIIA4-dTAG and GFP were transduced with normal or non-blocking (no base complementarity to the sgRNA scaffold) riboswitch fused sgRNAs targeting *GFP* or an *AAVS1* targeting control sequence. Cells were treated with increasing amounts of theophylline (0, 0.1, 0.5, and 1 mM) and DMSO or 200 nM dTAG-13. Loss of GFP fluorescence measured by flow cytometry 7 days after treatment. **B.** Repression of CRISPR activity via 16-nt riboswitch and AcrIIA4-dTAG. HCECs stably expressing Cas9 with AcrIIA4-dTAG and GFP were transduced with normal or 16-nt blocking riboswitch fused sgRNAs targeting *GFP* or an *AAVS1* targeting control sequence. Cells were treated as in Fig. **S2A**, and GFP fluorescence determined by flow cytometry.

**Figure S3.**
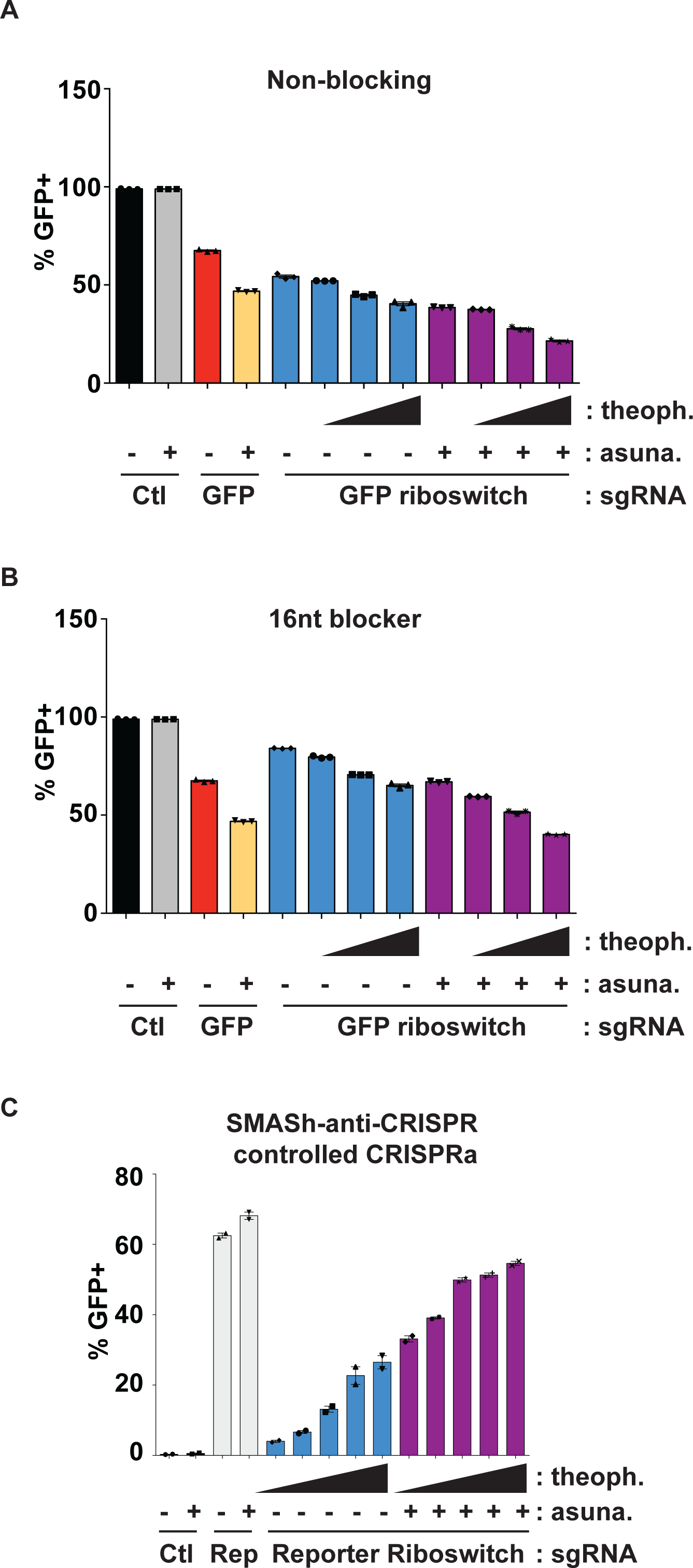
SMASh-tagged AcrIIA4 and riboswitch combination. **A.** A non-blocking riboswitch does not repress CRISPR activity. HCECs stably expressing Cas9 with SMASh-AcrIIA4 and GFP were transduced with normal or non-blocking (no base complementarity to the sgRNA scaffold) riboswitch fused sgRNAs targeting *GFP* or an *AAVS1* targeting control sequence. Cells were treated with increasing amounts of theophylline (0, 0.1, 0.5, and 1 mM) and DMSO or 1 μM asunaprevir. Loss of GFP fluorescence measured by flow cytometry 7 days after treatment. **B.** Repression of CRISPR activity via 16-nt riboswitch and SMASh-AcrIIA4. HCECs stably expressing Cas9 with SMASh-AcrIIA4 and GFP were transduced with normal or 16-nt blocking riboswitch fused sgRNAs targeting *GFP* or an *AAVS1* targeting control sequence. Cells were treated as in Fig. **S3A**, and GFP fluorescence was measured by flow cytometry 7 days after treatment. **C.** Elimination of leakiness by combining SMASh-AcrIIA4 with riboswitch controlled sgRNAs. HCECs stably expressing dCas9-VPR with SMASh-AcrIIA4 and a GFP transgene controlled by a minimal SCP2 promoter were transduced with normal or riboswitch sgRNAs targeting the reporter promoter (sgReporter) or an *AAVS1* targeting control (Ctl). Cells were treated with increasing amounts of theophylline (0, 0.1, 0.5, and 1 mM) and DMSO or 1 μm asunaprevir for 7 days. GFP fluorescence was examined by flow cytometry.

## REFERENCES

1. Kleinstiver B.P., et al. Nature. 529, 490–495 (2016).

2. Dow L.E., et al. Nature Biotechnology. 33, 390–394 (2015).

3. Wang T., Wei J.J., Sabatini D.M., Lander E.S. Science. 343, 80–84 (2014).

4. Nihongaki Y., Kawano F., Nakajima T., Sato M. Nature Biotechnology. 33, 755–760 (2015).

5. Zetsche B., Volz S.E., Zhang F. Nature Biotechnology. 33, 139–142 (2015).

6. Bondy-Denomy J., Pawluk A., Maxwell K.L., Davidson A.R. Nature. 493, 429–432 (2013).

7. Hwang S., Maxwell K.L. CRISPR J. 2, 23–30 (2019).

8. Shen J., et al. Science Advances. 3, 1701620 (2017).

9. Nabet B., et al. Nature Chemical Biology. 14, 431–444 (2018).

10. Shalem O., et al. Science. 343, 84–87 (2014).

11. Horlbeck M.A., et al. Elife. 5, 19760 (2016).

12. Chavez A., et al. Nature Methods. 12, 326–328 (2015).

13. Harper J.W., Adami G.R., Wei N., Keyomarsi K., Elledge S.J. Cell. 75, 805–816 (1993).

14. Chung H.K., et al. Nature Chemical Biology. 11, 713–720 (2015).

15. Cheng H. et al. Molecular Biosystems. 12, 3370–3376 (2016).

16. Nakamura M., et al. Nature Communications. 10, 194 (2019).

17. Iwasaki R.S. et al. Nature Communications. 11, 1394 (2020).

18. Kundert K. et al. Nature Communications. 10, 2127 (2019).

19. Roig A.I. et al. Gastroenterology. 138, 1012–1021 (2010).

20. Sack L.M., et al. Cell. 173, 499–514 (2018).

21. Erb M.A., et al. Nature. 543, 270–274 (2017).

