## Supplemental DNA information for "Proteasomal control of anti-CRISPRs for the regulation of CRISPR/Cas9 activity using Cas9-ACROBAT"

**Supplemental DNA Files**

**Cas9 vectors with AcrIIA4-dTAG fusions**

1. pENTR AcrIIA4-dTAG-T2A-Cas9.dna – For CRISPR cutting
2. pENTR AcrIIA4-dTAG-T2A-dCas9VPR.dna – For CRISPRa
3. pENTR AcrIIA4-dTAG-T2A-dCas9-KRAB.dna – For CRISPRi

**Cas9 vectors with SMASh-AcrIIA4 fusions**

1. pENTR SMASH-AcrIIA4-2xHA-T2A-3xFLAG-Cas9.dna – For CRISPR cutting
2. pENTR SMASH-AcrIIA4-2xHA-T2A-dCas9-VPR.dna - For CRISPRa
3. pENTR SMASH-AcrIIA4-2xHA-T2A-3xFLAG-dCas9-KRAB.dna – For CRISPRi

**Fluorescent CRISPRa reporters**

1. pHAGE minCMV dTomato CRISPRa reporter.dna
2. pHAGE-4xTgR-SCP2-EGFP-neo CRISPRa GFP reporter.dna

**Riboswitch sequences**

1. sgReporter #5 16nt blocker riboswitch U6fwd.dna
2. sgReporter #6 16nt non-blocker riboswitch U6fwd.dna
3. sgReporter #7-2 19nt non-blocker riboswitch U6fwd.dna
4. sgReporter non-riboswitch U6fwd.dna

**Spreadsheet with sgRNA sequences used**

1. sgRNA sequences used.xlsx
